# Hybrid Breakdown in Male Reproduction Between Recently-Diverged *Drosophila melanogaster* Populations Has a Complex and Variable Genetic Architecture

**DOI:** 10.1101/2022.10.13.512131

**Authors:** Matthew J. Lollar, Timothy J. Biewer-Heisler, Clarice E. Danen, John E. Pool

**Affiliations:** Laboratory of Genetics, University of Wisconsin-Madison, Madison, WI, 53706, United States

**Author notes:** Department of Biology, Indiana University, Bloomington, IN, 47405, United States. PreventionGenetics, Marshfield, WI, 54449, United States.

**Keywords:** Hybrid breakdown, reproductive isolation, postzygotic isolation, genetic incompatibilities, male reproduction, *Drosophila melanogaster*, genetic variation, polymorphism

## Abstract

Populations no longer experiencing a sufficient rate of gene flow will accumulate genetic differences over time. One potential consequence of divergence between natural populations is hybrid breakdown, which can occur during secondary contact when untested allelic combinations in hybrids beyond the F1 generation are maladaptive and restrict gene flow. Hybrid breakdown is an important process in the development and maintenance of species boundaries, and has largely been studied between populations that are completely or nearly completely isolated. Here, we leverage the recent worldwide expansion of *Drosophila melanogaster* to investigate signatures of hybrid breakdown between populations that diverged within approximately the last 13,000 years. We did not find clear evidence for hybrid breakdown in viability or female reproductive performance. In contrast, we found that many but not all between-population crosses yielded an elevated fraction of second generation male offspring that were unable to reproduce. The frequency of non-reproducing F2 males varied among different crosses involving the same southern African and European populations, as did the qualitative effect of cross direction, implying a genetically variable basis of hybrid breakdown and a role for uniparentally inherited factors. The levels of male reproductive failure observed in F2 hybrids were not recapitulated in backcrossed individuals, suggesting the existence of incompatibilities with at least three partners. These results suggest that some of the very first steps toward reproductive isolation may involve incompatibilities with complex and variable genetic architectures, and they support the prediction that hybrid breakdown affects the heterogametic sex first. Collectively, our findings on polymorphic incompatibilities within *D. melanogaster* emphasize this system’s potential for future studies on the genetic and organismal basis of early-stage reproductive isolation.

**IMPACT SUMMARY:** The biological diversity that exists around the world is an emergent property of the generation of forms, which are commonly grouped into units we call species. The rate at which new species form can be influenced by the evolution of reproductive isolation, the inability of groups to interbreed. When reproductive isolation is studied in its nascent stages, researchers can gain critical insights into the genetic architectures and evolutionary forces underlying the earliest steps toward speciation. One process that may contribute to early-stage reproductive isolation is hybrid breakdown, when genetic incompatibilities in the offspring of hybrid individuals reduce their fitness. Here, we illuminate a complex pattern of hybrid breakdown among natural populations of *Drosophila* flies that diverged within the past 13,000 years. We find signals of hybrid breakdown involving male reproduction, between some but not all population pairs, whereas we find no clear evidence for hybrid breakdown impacting female reproduction or developmental survival. These findings are in agreement with Haldane’s Rule, which posits that hybrid incompatibilities are more likely to affect the sex that carries distinct sex chromosomes (here, XY males). From certain crosses between African and European fly strains, we find strongly elevated rates of reproductive failure in second generation hybrid males, but outcomes vary dramatically depending on the individual strains crossed. We also provide evidence of incompatibilities underlying male reproductive failure that involve three or more genes, including uniparental factors such as the Y chromosome or mitochondrial genome. Our results highlight a complex and variable basis of hybrid breakdown during the earliest stages of reproductive isolation, in contrast to commonly envisioned scenarios that focus on two-locus incompatibilities caused by fixed genetic differences between groups. These findings also suggest that recently diverged populations of *D. melanogaster* provide notable opportunities for future studies of the genetic basis of early-stage reproductive isolation.

## INTRODUCTION

Populations with limited genetic exchange experience distinct population genetic forces that may drive allele frequency change between groups through successive generations. As a consequence of this genetic divergence, reproductively isolated populations descending from a once freely interbreeding common ancestor may be partially or wholly reproductively isolated upon secondary contact (Mayr 1942). Alleles that are at appreciable frequency in one but not both populations are tested against a novel genetic background upon hybridization. Neutral or beneficial alleles in one population may prove deleterious in a hybrid genome, and consequent reductions to fitness may prevent reproduction or survivability in hybrids (Coyne and Orr 2004).

These interactions are ‘unseen’ by selection until the point of hybridization, and thus populations need not ever pass through unfit allelic intermediates for alleles involved in incompatibilities to evolve between them. Rather, they may evolve simply as a pleiotropic byproduct of independent evolution (Presgraves 2010). Collectively, these genetic incompatibilities were described by and attributed to Bateson (1909), Dobzhansky (1934) and Muller (1942) (BDM-incompatibilities or BDMIs). BDMIs are relevant to many species definitions, particularly those which generally define species boundaries by the presence of strong or complete reproductive isolation between groups (Coyne and Orr 2004).

The frequent existence of BDMIs between species has been well-documented by field and lab studies (Blackman 2016), and found in both systems that are partially or completely isolated. Analysis of studies characterizing reproductive isolation have revealed some general patterns. The primary and most consistent pattern to emerge within taxa such as *Drosophila* is that the strength of reproductive isolation is positively correlated to genetic distance (Orr 2005). As all genes act within the epistatic context of their genetic background, it is perhaps expected that the number of possible BDMIs between genomes is elevated with increasing differences between them (Satokangas *et al*. 2020). Theoretically, incompatibilities are expected to accumulate at a rate proportional to the square number of differences between populations (Orr and Turreli 2001, Matute *et al*. 2010).

While between-species models have been successful in identifying genomic regions and sometimes genes underlying hybrid dysfunction (Blackman 2016), a remaining challenge in these studies is the determination of which incompatibilities have historical relevance to the speciation process. Inherent to the study of distantly-related species is the assumption that detectable incompatibilities in the present were involved in the generation of initial barriers to gene flow. The accumulation of incompatibilities over time may strengthen initial barriers to hybridization or even ‘complete’ the speciation process (Dobzhansky 1937), but there is no certainty that currently-known BDMIs were involved in the early stages of isolation (Coyne and Orr 2004).

Many of the outstanding questions remaining in the field of speciation are difficult to answer in between-species studies. The determination of which evolutionary forces govern the fixation or rise in frequency of incompatibility variants may not always be possible for well-isolated species, as any population genetic signatures of positive selection may no longer be detectable. In addition, if there was a polymorphic basis to early-acting incompatibilities between diverging taxa (Cutter 2012), this history may be obscured when incompatible variants eventually become fixed. Although polymorphisms contributing to incompatibilities may often be present in the early stages of reproductive isolation (Reed *et al*. 2008, Cutter 2012; Laturney and Moehring 2012), complete isolation inevitably involves the fixation of alleles between populations (Larson *et al*. 2018), with the potential to obscure a history of polymorphic incompatibility. Furthermore, the genetic tools available to pursue the basis and mechanism of incompatibilities between completely-isolated taxa are inherently limited, especially if such incompatibilities between species result in lethality of hybrids (Presgraves 2007). Researchers have used incompletely-isolated species models to overcome limitations to studies in distantly related species, and to determine whether the speciation process differs at distinct evolutionary stages (Kulmuni *et al*. 2020). Taking this approach to the extreme, if studies of very recently diverged populations within the same species can identify incompatibilities that are present at some polymorphic frequency, new insights into the biology of emergent reproductive isolation may be elucidated.

Although populations that have experienced a recent divergence may be separated genetically only by a small number of alleles, evidence for BDMI-like incompatibilities between incompletely-isolated populations within species has been documented (Coughlan and Matute 2020). In such cases alleles involved in an incompatibility may or may not be fixed within a population (Turelli and Orr 2000), and if they are recessive, isolation will typically only manifest in second generation or later hybrids of sexually reproducing diploids. Reproductive isolation of this kind is often termed “hybrid breakdown” and is a specific category of intrinsic post-zygotic reproductive isolation (Oka *et al*. 2003).

Investigation of hybrid breakdown and later generation BDMIs has received some attention in recent speciation literature, and is often pursued between closely-related species (Breeuwer and Werren 1995, Oka *et al*. 2003, Renaut and Bernatchez 2011, Matsubara 2020, Morgan *et al*. 2020). Examples of within-species models of reproductive isolation are more limited (White *et al*. 2012, Pritchard and Edmands 2013, Stelkens *et al*. 2015, Koski *et al*. 2022), yet in these few studies the existence of moderately strong isolating barriers among populations has been evidenced. Still, there remain gaps in our understanding of the biology underlying the earliest incompatibilities segregating within species, the evolutionary forces that act upon causative agents, and whether hybrid breakdown contributes significantly to reproductive isolation in its nascent stages. Within-species allopatric populations are particularly fertile territory for the investigation of such questions during early-stage reproductive isolation, and one potential candidate for such studies is the vinegar fly *Drosophila melanogaster*, a species with many recently-isolated populations and the most intensively-studied species in a genus with broad importance in the speciation literature (Coyne and Orr 1989).

Natural populations of *Drosophila melanogaster* are thought to have originated in sub-Saharan Africa, where at some unknown time it formed a commensal relationship with humans (Lachaise *et al*. 1988; Lachaise and Silvain 2004). It is thought that expansion out of a southern-central African ancestral range (Pool *et al*. 2012) first occurred approximately 13,000 years ago (Sprengelmeyer *et al*. 2020). While this period of population divergence encompasses nearly 200,000 fly generations, it represents only a brief interval of population genetic time (less than 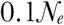 generations, based on the estimated long term ancestral population size; Sprengelmeyer *et al*. 2020). The species then expanded across much of sub-Saharan Africa, and soon thereafter crossed what is now the Saharan desert, which involved a moderately strong population bottleneck leading to reduced genetic diversity outside of sub-Saharan Africa (Begun and Aquadro 1993; Sprengelmeyer *et al*. 2020). However, the species may have only entered Europe roughly 1,800 years ago (Sprengelmeyer *et al*. 2020; Figure 1D), and it was not observed in northern Europe until the 1800s (Keller 2007). Additionally, phenotypic differences have evolved that correlate with geography, providing evidence of local adaptation between populations within and beyond Africa (*e.g*. David and Capy 1988; Pool and Aquadro 2007; Fabian *et al*. 2015; Lack *et al*. 2016a). Populations of *D. melanogaster* are now found worldwide, providing a rich pool of genetic and phenotypic diversity that can be readily studied in the laboratory (*e.g*. Lack *et al*. 2016b).

**Figure 1.**
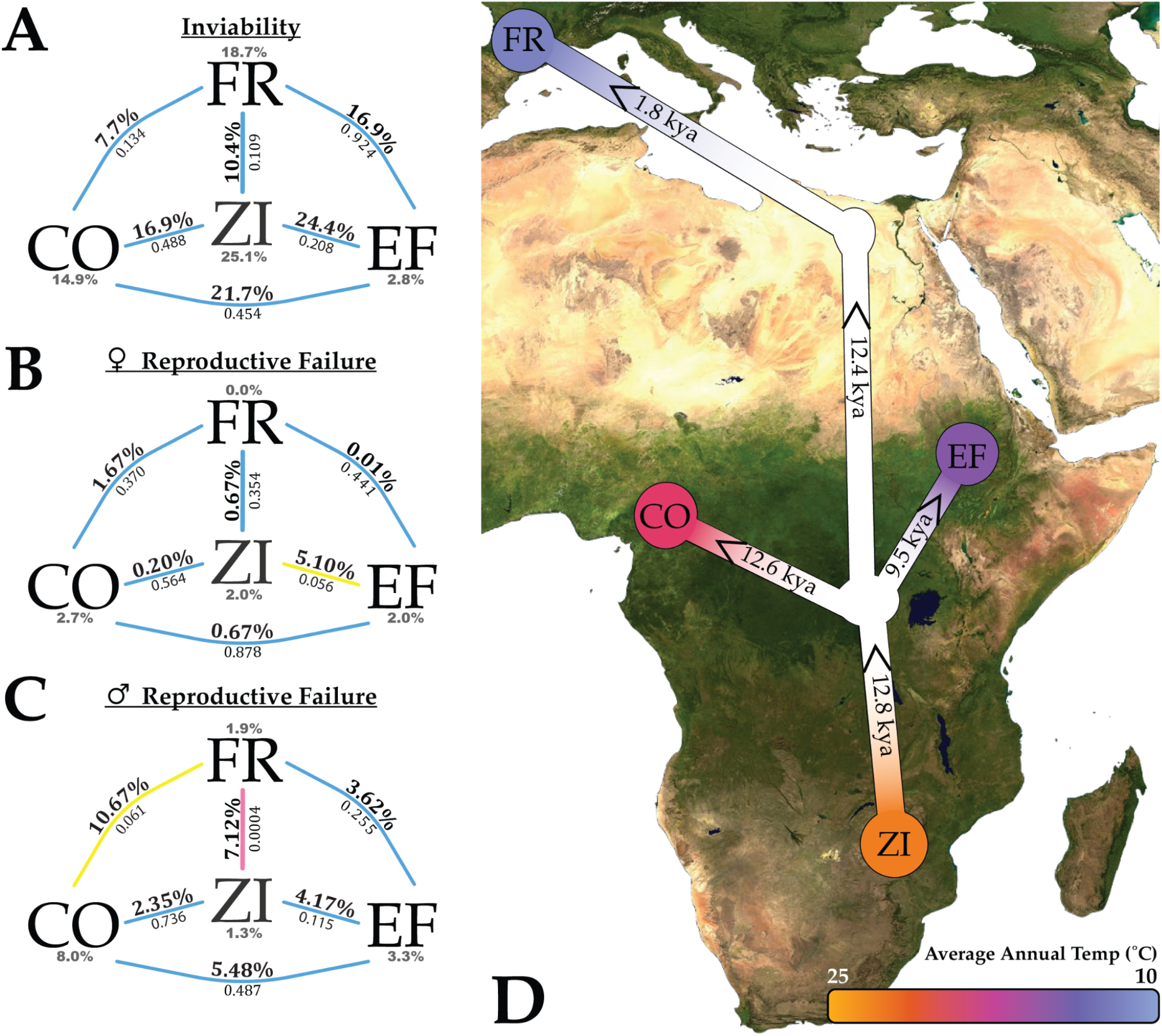
Failure in male reproductive success is the predominant mode of hybrid breakdown among F2 hybrids between various *D.melanogaster* populations. Left: F2 combined cross values and significance for average percent inviability (**A**), and for female (**B**) and male (**C**) reproductive failure rates. Line color denotes significance of differences in inviability/reproductive failure between population’s crosses relative to both within-population cross groups, where pink indicates p<0.05, yellow indicates 0.05<p<0.1, and blue indicates p>0.1. Within-population cross averages (gray) are listed adjacent to the population label. P-values were generated from bootstrapping (see Materials and Methods) and are listed underneath each between-population percentage. Individual cross averages are supplied in Tables S1-S3. **D**: Map of *D. melanogaster* population expansion and estimated divergence times from Sprengelmeyer *et al*. (2020). Divergence is given in thousands of years (kya) and each value represents the initial expansion estimate between nodes. Expansion out of ancestral ranges in southern-central African occurred roughly 12.8 kya. Populations rapidly colonized west Africa (12.6 kya), and later other regions of central Africa including our Ethiopian sample (9.5 kya). An additional estimated split (2.7 kya) between low and high altitude Ethiopian populations, the latter being used in the present study, is not depicted. Trans-Saharan migration occurred roughly 12.4 kya, during which populations experienced a moderately strong population bottleneck. Migration from the Middle East into cooler European regions occurred in the more recent past (1.8 kya). Coloration based on average annual temperature (scale bottom right) highlights the dramatic environmental differences between regions.

Evidence of partial reproductive isolation between populations within the *D.melanogaster* species has previously been suggested, particularly between tropical African and temperate non-African populations. Prezygotic barriers have been reported, evidenced primarily by the unidirectional mating preference of Zimbabwe-type females, who disfavor males of cosmopolitan origin in favor of other ‘Z-type’ males (Hollocher *et al*. 1997, Grillet *et al*. 2011). A separate instance of unidirectional mating preference has also been reported between populations of western/central African origin versus all others (Bontonou *et al*. 2013). Post-zygotic isolation after the F1 generation has also been reported within the species. When X chromosomes from North American strains of high European ancestry were introgressed into autosomal backgrounds of Caribbean strains with larger African ancestry, lethality was observed in over half of all crosses, and 57% of the viable crosses showed male and/or female sterility. The reciprocal introgression produced F2 lethality only in 18% of crosses, and sterility in around 30% of the viable crosses (Lachance and True 2010). Reduced egg-to-adult viability has also been observed in some crosses between African and non-African strains (Alipaz *et al*. 2001) and from naturally admixed populations with more even levels of African and European ancestry (Kao *et al*. 2015). Thus, there is good evidence that incompatibilities may be exposed in crosses between pure African and European strains.

Subsequent genomic inference of pervasive epistasis in admixed *D. melanogaster* (Pool 2015, Corbett-Detig *et al*. 2013) further bolster the desire to pursue the *D.melanogaster* system as a model for early-stage reproductive isolation. Pool (2015) assessed European/African ancestry across genomes of the *Drosophila melanogaster* Genetic Reference Panel (Mackay *et al*. 2012) and found strong evidence for natural selection mediating the outcome of ancestry within strains. Particularly, there was a genome-wide signal for ancestry disequilibrium between unlinked loci in these highly-inbred lines (Pool 2015). Ancestry disequilibrium represents the deficiency of Africa-Europe allele combinations between locus pairs that is not predicted under neutrality when accounting for the number of admixed generations. Results found here suggest that natural selection and negative epistasis mediated the outcomes of natural admixture between European and African alleles and laboratory inbreeding. Additionally, epistatic incompatibilities were inferred to segregate between natural populations of *D.melanogaster* (Corbett-Detig *et al*. 2013), based on the genomes of laboratory-admixed *Drosophila* Synthetic Resource Panel strains (King *et al*. 2012). Taken together, these studies highlight the possibility of using genomic variation and laboratory experiments to detect signatures of isolation within the species.

Here, we sought to investigate the presence of hybrid breakdown among several natural populations of *D. melanogaster*. We find little evidence of breakdown for inviability or hybrid female reproductive failure, but provide substantial evidence for hybrid male reproductive failure in the F2 generations, especially from crosses between France and Zambia populations. We find that both frequencies of non-reproductive males and the effect of cross direction are highly dependent on the specific strains crossed. Additionally, we find that backcrosses among crosses with male reproductive failure do not recapitulate or enrich the phenotype of F2 crosses, suggesting that incompatibilities underlying reproductive failure do not conform to two-locus BDMI models. Our investigation suggests that early-stage reproductive isolation within the species has a variable and complex genetic architecture, and that *D. melanogaster* represents a promising system for studying the genetic basis of early-stage reproductive isolation.

## MATERIALS AND METHODS

### Fly husbandry

All stocks and crosses were reared in bottles containing standard *Drosophila* cornmeal/molasses medium, prepared in batches consisting of 4.5 L water, 500 mL cornmeal (Quaker yellow), 500 mL molasses (Grandma’s unsulphured), 200 mL powdered yeast (MP Biochemical Brewer’s), 54 g agar (Genesee *Drosophila* type II), 20 mL propionic acid, and 45 mL 10% Tegosept solution in 95% ethanol (by volume). Bottles were maintained at a constant 25°C temperature and 12 hour light/dark circadian cycles. Humidity was not strictly controlled, but typically ranges between 50-60%. Bottles used for virgin collections were moved to an 18°C environment during hatching periods, and collections were maintained under the same conditions as stocks. Crosses were founded using approximately 10-20 flies per strain, and F1 interbreeding occurred using an estimated 50-100 F1 flies per bottle. Flies were transferred to fresh bottles every three days, and offspring were collected from crosses up to 16 days from the first transfer of flies to the bottle. F2 offspring in all assays were collected 24-48 hours post-eclosion and placed into vials containing no more than 20 flies per vial. Flies were aged for an additional 24-48 hours before use in assays. Virgin females used in male no-choice mating assays were collected into vials of no more than 20 females per vial, and aged 2-5 days before use in assays.

### Generation of experimental individuals

We first selected six unique wild-derived inbred strains from each of four natural populations, representing Cameroon, France, high-altitude Ethiopia, and Zambia sources (Pool *et al*. 2012). In all cases except for Cameroon, strains free from commonly known inversions were selected, while all available inbred strains from our Cameroon collection contain at least one known inversion. Six between-population crosses were performed between each combination of population pairs, for a total of six population pairings (Tables S1-S3). Each of the six crosses between two populations involved different strains, and three were performed in each cross direction. In crosses involving Cameroon, one Cameroon strain was used for two crosses, as we possessed five inbred strains from this sample. Three within-population crosses for each of the four source populations were also performed and used as controls for statistical comparison. First generation (F1) hybrids of crosses were subsequently intercrossed to generate second generation hybrids (F2). Reproductive isolation in F2 hybrids was then measured by three metrics: Egg to adult viability and reproductive success in no choice mating success assays for both males and females.

For the larger-scale male reproduction screen between France and Zambia, we randomly selected five inversion-free strains each from both population samples, and this time we performed all pairwise cross combinations among these strains. Each of the 25 cross pairs were sampled in both cross directions for a total of 50 between-population crosses. Within-population controls were likewise generated through pairwise crosses among each of the five strains within a population, crossed in both directions, yielding 40 within-population control crosses consisting of 20 crosses each within France and Zambia (Tables S4 & S5). F2 males were generated and assayed identically to the methods described above for the four-population analysis.

Backcrossed (BC) males were generated by two primary cross designs to test for effects of either the mitochondria or Y chromosome. For tests involving a focal Y chromosome, F1 males were crossed back to F0 females from the non-focal ancestral type (thus pairing the focal Y chromosome genotype with homozygous autosomal and/or X chromosomal genotypes in many backcross offspring). For tests involving a focal mitochondrial genotype, F1 females were crossed back to F0 males from the non-focal ancestral type (thus pairing the focal mitochondrial genotype with homozygous autosomal and/or X chromosomal genotypes in many backcross offspring). Male reproductive success in BC males was measured in an identical procedure as for the F2 studies above.

### No-choice mating reproductive success assay

F2 males were paired with two virgin females (one from each population involved in the cross) in a single vial, and allowed to mate for nine days. Females used in the assay were randomly chosen from all employed lines for a population, excluding those that founded the cross being assayed. After nine days, vials were checked for the presence of larvae or pupae. If no offspring were present, males were transferred to a new vial, paired with two new virgin females, and allowed an additional nine days for possible mating. Males that were successful in the first trial were not assayed in a second trial and considered successful. Males that failed to produce offspring in the second 9 day trial were counted as non-reproducing, and successful males in either trial were counted as reproducing. If a male or both females were deceased during either checkpoint, results from that male were discarded. Assays were conducted in ambient lab conditions.

Female reproductive performance assays were conducted identically to male assays, with the exception of pairing a single hybrid female to two male mates, again randomly selected from strains of each population represented in the cross but excluding the strain used to establish a cross. Reproductive success was counted as a binary trait (fecundity was not assayed quantitatively).

### Egg-to-adult viability assays

F1 offspring were collected into vials of no more than 25 flies per vial at 0-48 hours post-eclosion. Flies were aged an additional 3-5 days in cohorts of vials containing both males and females. At the time of assay flies were separated in groups of three males and three females, and placed into fresh vials to allow mate/laying for an additional 18-22 hours (density of eggs in vials was not strictly controlled). At this time, flies were discarded and all embryos were counted. Larvae were added to the egg count if embryos had hatched before counting occurred. Vials were maintained at 25°C for an additional 15-17 days, at which point adults were counted from vials. Trials with less than five embryos counted were not considered. Unless otherwise noted in Table S1, 10 trials per cross were performed for each cross.

### Significance testing

We sought to test whether the levels of reproductive failure observed in each between-population test cross exceeded our null expectations based on the levels of reproductive failure observed in within-population control crosses. P-values for all reproductive failure analyses were generated using a bootstrap resampling approach to compare a given between-population cross result against our full panel of within-population control crosses from the same two populations. For each of one million bootstrap replicates, a random control cross was first selected from those obtained from both parental populations. Second, the control individuals were resampled with replacement, based on the control cross’s own sample size, to account for sampling variance in the control cross data. Third, we again resampled individuals from the (resampled) control cross data, this time based on the between-population test cross’s sample size, to account for sampling variance in the test cross data. Fourth, if the number of reproductive failures after the latter resampling step was greater than or equal to the number observed in the test cross, then this replicate was added to the numerator of the p-value calculation (which had a denominator equal to the number of replicates, one million). P-values therefore represented the proportion of replicates in which our two-step resampling of control cross data yielded at least as many reproductive failures as observed for the test cross.

When we instead sought to test whether a full set of crosses between two populations departed from the null expectations of within-population crosses, we followed a similar procedure. Here in each replicate, we chose a random control cross for each test cross (sampling control crosses with replacement), followed by the same two-step resampling scheme as described above, and asked whether the total number of reproductive failures (summed across these resampled crosses) was greater than the total actually observed across all of the between-population crosses being considered.

For our four population analysis, we only tested whether the full set of crosses between two populations was significantly different than the panel of within-population crosses from the same two populations in a pair. In the case of the larger France/Zambia screen for male reproductive failure (Figure 2; Tables S2 & S3), where we also tested the significance of individual crosses, a Benjamini-Hochberg multiple test correction was applied to these p-values using the p.adjust method “BH” (Benjamini and Hochberg 1995) from the Rstats package (version 3.6.2)(R core 2021). For this same France/Zambia comparison, we also tested whether individual between-population crosses showed significant differences between the reproductive failure rates observed between the two cross directions. Here, the cross direction with the lowest rate of reproductive failure was considered the control set, and tests were performed identically to the description above for testing a full set of crosses between two populations.

**Figure 2.**
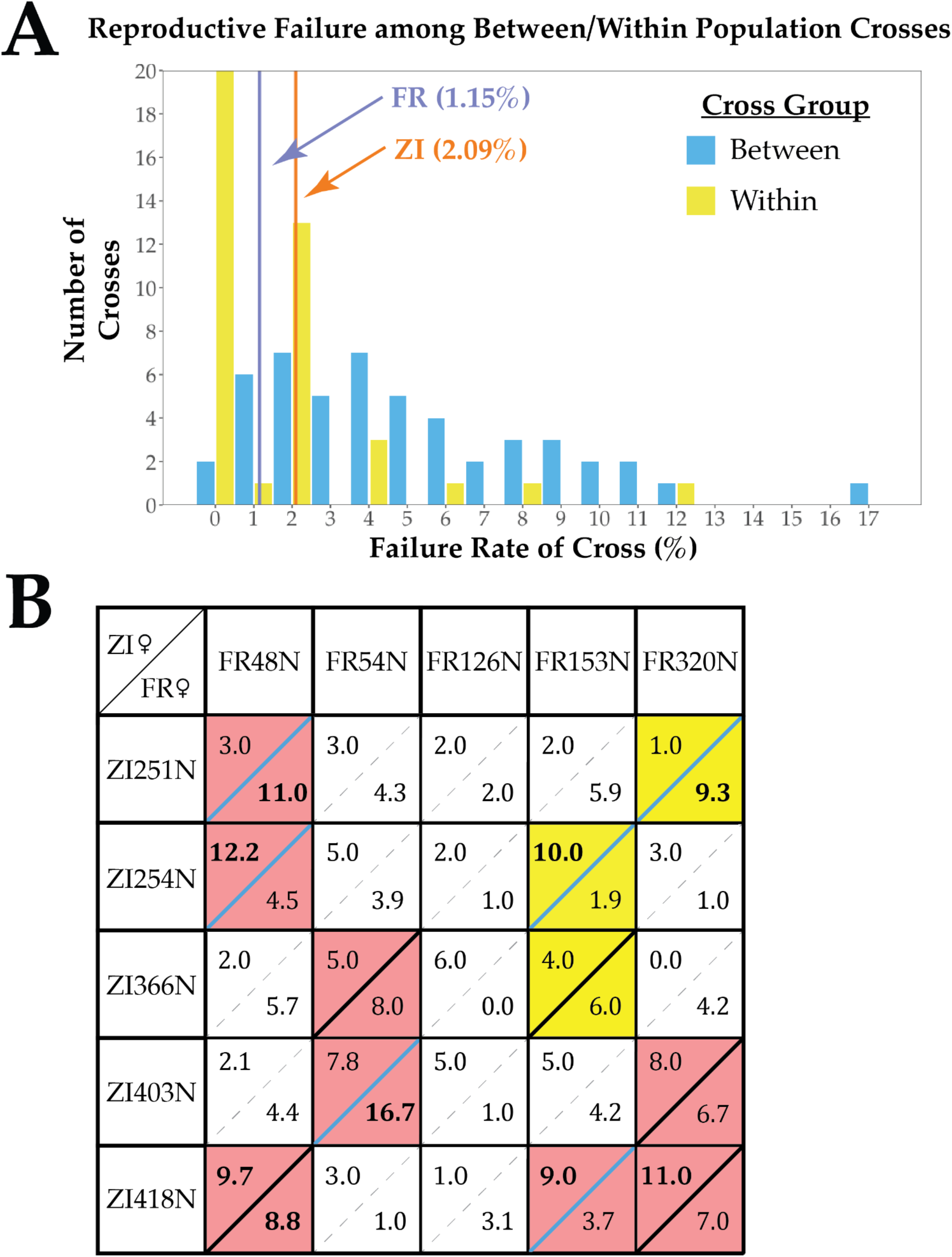
Male reproductive failure rates between French and Zambian *D.melanogaster* crosses deviate significantly from within-population crosses, and have a variable and sometimes crossdirection-specific pattern. **A:** Histogram depicting percentage of male reproductive failure for between-population (blue) and within-population (yellow) cross directions. Values on the x-axis are the percentage of males that failed in all four reproductive assays, each involving one female from each parental population (binned by rounding percentages up to nearest whole number). Within-population cross averages are displayed for France (blue line) and Zambia (orange line) cohorts. In total, 36 of the 50 between population crosses have reproductive failure rates higher than both within-population averages (bins right of both lines), and the rate of reproductive failure between these cohorts is significantly different (bootstrap p<1.0× 10^-6^). Individual cross counts are supplied in Tables S4 & S5. **B:** Male F2 reproductive failure rates among all possible crosses between five France and five Zambia strains, for both cross directions. Each cross is represented by a separate male failure rate for each cross direction (FR maternal top-left, ZI maternal bottom-right). Individual cross directions with significant failure rates compared to within-population controls are bolded. Crosses whose direction-combined cross failure rates have significant raw p-values (p<0.05) are highlighted pink, and those with 0.05<p<0.1 are highlighted yellow. These highlighted crosses were tested for differences between cross directions in failure rate, and crosses with significant (p<0.05) differences are denoted by a blue solid line.

One-Way ANOVA was used to analyze viability data. ANOVAs were conducted in R. Full code and documentation of the above methods can be found at [www.github.com/mjlollar/significance_testing].

## RESULTS

We first searched for evidence of hybrid breakdown involving three traits (egg-to-adult viability, female reproductive ability, and male reproductive ability) among four natural populations of *D. melanogaster* (Cameroon, high altitude Ethiopia, France, and Zambia). For each population pair, we performed six unidirectional crosses involving separate inbred strains - three with each population as the maternal parent. We applied assays described in the Materials and Methods section to F2 individuals. We then tested for differences in survival/reproduction between the sets of between-population crosses and within-population crosses from a given population pair, in order to assess potential evidence for hybrid breakdown between *D. melanogaster* populations.

We did not find clear evidence of egg-to-adult viability decline between any population pair when compared to within population cohorts (Figure 1A). In general, embryo yield varied both between crosses and within crosses, and assay conditions may have impacted our ability to detect subtle inviability differences. Populations that laid many eggs may have suffered from overcrowding affecting the hatch rate of an assay replicate, diminishing the population average viability rate. This may be evidenced, for instance, by the Ethiopian population (Table S1). Previous studies in our lab have found lower egg lay rates in this cohort (Lack *et al*. 2016c), and indeed the egg counts tended lower in our assay (Table S1), while overall inviability within the Ethiopian population was much lower than other within-population samples. We note that alternative assays could reveal more subtle viability differences between populations, but our screens do not suggest that viability under benign lab conditions is strongly impacted by hybrid breakdown between *D. melanogaster* populations.

Female reproductive ability also did not significantly differ between population crosses compared to within-population samples (Figure 1B). In general, failures in the mating assays performed among all cohorts were almost entirely absent (Table S2). A marginally significant signal between France and Ethiopia (bootstrap p=0.0558) was driven primarily by a single between-population cross out of six with a high rate of failure (Table S2). We do not interpret this result as clear evidence of hybrid breakdown, although we can not rule the existence of polymorphic incompatibilities between these populations affecting this trait. We also note that our study did not examine quantitative female fecundity, which might conceivably be affected by hybrid breakdown.

In contrast to the above traits, we find two signatures of increased F2 male reproductive failure between populations: a strong signal between France and Zambia (Figure 1C; bootstrap p=3.76×10-4) and a weaker signal between France and Cameroon (Figure 1C; bootstrap p=0.06117). Interestingly, both pairs demonstrated a cross-direction pattern of reproductive failure (Table S3). Crosses involving France and Zambia were more often enriched for male failure in the Zambian maternal / France paternal direction of crosses (bootstrap p<1.0×10-6; Table S3), while crosses involving France and Cameroon failed more frequently in paternal Cameroon / maternal France direction (bootstrap p<1.0×10-6; Table S3). Despite the high average sterility rate between France and Cameroon, the statistical signal was diminished due to the presence of a single within-population control cross with notable high failure rate (Table S3) in the Cameroon population. While a larger sample may improve the statistical comparison for this population pair, we again note the difficulty in the immediate interpretation of this result, in part due to the presence of inversions among our assayed Cameroon strains. Conversely, the strong signal of male reproductive failure between France and Zambia cross pairings represented a more promising suggestion of hybrid breakdown, and prompted us to investigate this trend of elevated male reproductive failure in greater detail.

We next sought to increase overall sampling among France and Zambia crosses to further assess evidence of hybrid breakdown between these populations and to examine its genetic architecture. We chose 5 inbred strains from each population and performed all possible between-population and within-population crosses among them (making 50 total cross directions for between-population crosses and 40 within). Results of these experiments qualitatively supported our preliminary findings of elevated male reproductive failure from crosses between France and Zambia strains. Within-population crosses only yielded average male reproductive failure rates of 1.15% (France) and 2.09% (Zambia), and similarly we found among males sampled from within these ten inbred strains, an average of just 1.4% of failed to reproduce (Table S4). In contrast, crosses between these populations produced a mean male failure rate of 4.94%, more than triple the within-population average (Figure 2A; Table S5). And yet, the latter average masks striking variation among between-population crosses. Averaging between cross directions, five of the 25 between-population crosses gave male failure rates no greater than the within-Zambia average cited above. In contrast, a cross between FR54N and ZI403N gave an average of 12.2% male failure between the two cross directions, and several other crosses showed notable elevations as well (Figure 2B).

We applied a bootstrap resampling approach to compare individual between-population crosses against the panel of within-population crosses, to ask which of them showed evidence for elevated rates of F2 male reproductive failure. Nine of the 50 between-population cross directions were found to be significant at p<0.05 (Figure 2B, bold values). When both cross directions were considered as a single sample, eight crosses out of 25 were significant at p<0.05 (Figure 2B, pink boxes), while another three crosses were marginally significant at 0.05<p<0.1 (Figure 2B, yellow boxes). We applied a Benjamini-Hochberg multiple test correction to the resampling p-values (Benjamini & Hochberg 1995), and found that no individual cross could be strictly considered significant (Table S5). Despite this, 36 out of the 50 individual between-population crosses had a percent failure rate higher than both within-population averages (Figure 2A). When all between-population crosses were tested jointly, no replicate among 1,000,000 permutations matched the total observed number of failed F2 males. Collectively, these data suggest that male reproductive failure is significantly elevated in between-versus within-population crosses between France and Zambia *D. melanogaster*.

Finally, we tested the significance of cross direction in the rate of F2 male failure, as suggested from preliminary screening (Figure 1, Table S5). We considered all crosses with combined-direction p-values below 0.1, a relaxed threshold allowing us to test a total of 11 cross (Figure 2B, highlighted crosses). Roughly half of these crosses (6/11) showed significant differences between cross directions (Figure 2B, blue solid lines). Of these six, three crosses produced F2 failures more significantly in the France maternal cross direction, and three crosses more frequently in the Zambian maternal direction. The most extreme ratio between cross directions, from cross FR320N/ZI251N, involved 9.3% F2 male failure with a France mother, but only 1.0% with a Zambia mother. The greatest arithmetic difference in failure rates, for cross FR54N/ZI403N, involved 16.7% failure with a Zambia mother, versus 7.8% with a France mother. These results do not support the above preliminary finding of a France paternal / Zambia maternal pattern of male failure in the preliminary survey. Rather, they suggest a variable pattern to male reproductive failure that is sometimes influenced by parental sex in either direction.

The asymmetric cross direction pattern of F2 male reproductive failure observed in some crosses implies the involvement of uniparentally inherited chromosomes, *i.e*. Y chromosomes or mitochondria (unless other cytoplasmic factors are involved). One simple way to test for effects from these uniparental factors is to employ backcross designs aimed at exposing each uniparentally inherited unit to a homozygous (or hemizygous) genetic background of the alternate ancestry. If the potential underlying genetic basis of male reproductive failure follows a two-locus recessive BDMI model involving a uniparental chromosome, the expected failure rate among BC males should be higher in some backcrosses relative to F2 hybrids from a F1 intercross (Figure 3).

**Figure 3:**
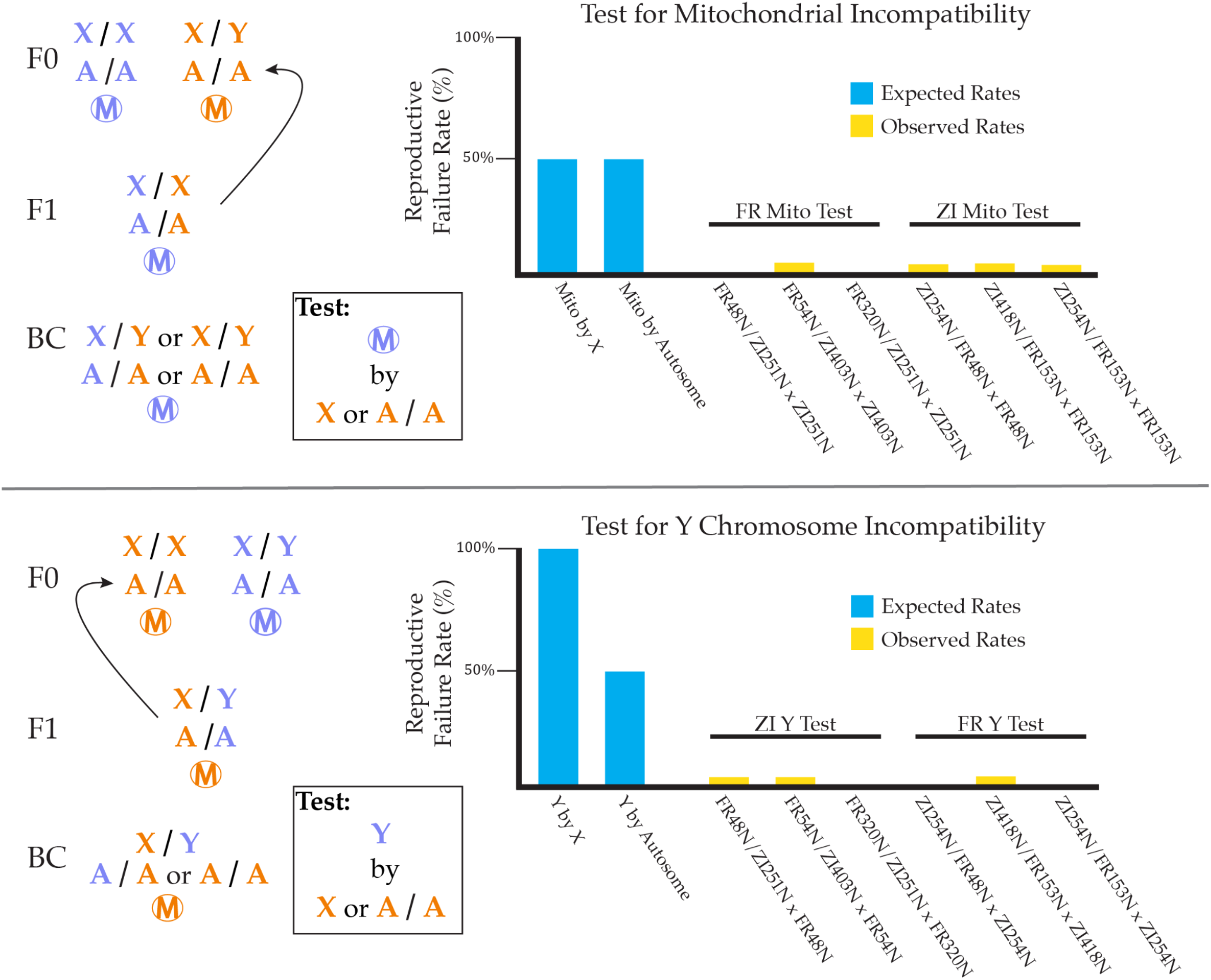
Male reproductive failure in backcrosses is reduced relative to F2 males, suggesting that a two-locus BDMI model is not appropriate for incompatibilities involving uniparental factors. Left: Backcross design to test for mitochondrial (top) or Y (bottom) by X/autosome two-locus recessive BDMIs between two populations (blue and orange). Above, an example cross design to test for BDMIs involving the blue mitochondrial haplotype. Below, an example design to test for BDMIs involving the blue Y haplotype. Right: Reproductive failure rates among BC males. Under the hypothesis of a single, two-locus recessive BDMI, these backcrosses increase the probability of uniparental by hemizygous X or recessive autosomal genotypes, and the expected rate of reproductive failure among backcrosses should increase (blue bars). Out of the six crosses in Figure 2B that had cross direction effects, three crosses each produced reproductive failure at higher rates in the France or Zambia maternal direction. This allowed us to test for BDMIs involving four uniparental elements (France or Zambia, mitochondria or Y), as indicated by labels above yellow bars. Backcrosses are written in the format of “(F1 Maternal Parent / F1 Paternal Parent) x F0 partner”, where the F0 partner is male for backcrosses targeting the mitochondria (top) and female for backcrosses targeting the Y chromosome (bottom). Individual cross counts are supplied in Table S6. Male reproductive failure was virtually absent in all six focal backcrosses assayed (yellow bars), differing drastically from expected values under two-locus BDMI models (blue bars). These results suggest that pairing a uniparental chromosome with hemizygous/homozygous genotypes from the opposite population is not sufficient to unmask the incompatibilities involved in cross direction asymmetry of F2 hybrid breakdown, indicating that these do not reflect classic two-locus BDMIs.

We performed two backcrosses on each of the six of the France/Zambia cross directions identified as having significantly greater male failure rates than their reciprocal crosses (Figure 2B) - one meant to expose a mitochondrial incompatibility, and the other meant to expose a Y chromosome incompatibility. We predicted that two-locus BDMIs involving a maternally- or paternally-inherited partner would yield 50% or more of males carrying incompatible genotypes in the relevant backcross, even if an autosomal or X-linked partner locus was recessive (Figure 3). In stark contrast to this prediction, all 12 backcrosses yielded male failure rates below 5% (Figure 3; Table S6). Rather than being sharply elevated, each of these male failure rates were lower than observed for F2 males from the enriched cross direction. These findings suggest that potential male reproduction BDMIs associated with uniparental chromosomes are unlikely to involve only a single BDMI partner. Instead, the reduced male failure rates from backcrosses could indicate that BDMIs involving uniparental chromosomes require at least one more partner locus from the same parental strain as the uniparental factor (which can not be homozygous under our backcross design), in addition to one or more partner loci from the opposite strain.

## DISCUSSION

The accumulation of reproductive isolation can occur as a consequence of restricted gene flow and independent evolution between populations. The strength of isolation has important consequences for the maintenance and origins of biological diversity (Rabosky 2016), and is considered to be a determining factor in species boundaries by many definitions (Coyne and Orr 2004). Even if isolation is incomplete and reproductive barriers are weak, incompatible alleles may be present between populations. When such variants are recessive, barriers to reproduction manifest in F2 or later generation hybrids, in a process commonly referred to as hybrid breakdown. Here, we provide evidence for hybrid breakdown in male reproduction between various natural populations of *D. melanogaster*. This study highlights the potential of using *D. melanogaster* as a powerful within-species model of early-stage reproductive isolation, and provides preliminary evidence that such incompatibilities may be more genetically complex in nature than what might be expected under the simplest BDMI models.

Our analysis of four *D. melanogaster* populations revealed that hybrid breakdown between populations was most clearly reflected in a decline in male reproductive success in the F2 generation, whereas inviability and female reproductive failure occurred at relatively similar rates in between- and within-population crosses. In contrast to our largely negative results for the latter traits, Lachance and True (2010) did find evidence for inviability and female sterility, in addition to male sterility, when they created strains that were homozygous for X chromosomes and autosomes from contrasting geographic origins (e.g. crossing an X chromosome from a majority African population into an autosomal background of primarily European ancestry). The fully homozygous nature of those X-autosome combinations may have given that study a greater ability to detect some incompatibilities. However, the crossing scheme employed in that study also leaves the possibility that female sterility might have been caused by simple recessive deleterious X-linked variants (whereas recessive X-linked inviability and male sterility variants are less frequently encountered during lab inbreeding, due to more efficient natural selection against less-fit hemizygous males). Additional evidence for inviability incompatibilities comes from the studies of Alipaz *et al*. (2001) and Kao *et al*. (2015). It is possible that further F2 studies like ours may yield further evidence for hybrid breakdown involving viability and female fertility, whether by expanding the experimental scale, controlling for population differences in egg production, quantifying female fecundity, or performing assays under more challenging environmental conditions. Still, on its own, our current study provides no clear support for hybrid breakdown between *D. melanogaster* populations leading to inviability or female reproductive failure.

Our deeper examination of hybrid breakdown in male reproduction between populations from France and Zambia revealed a strikingly variable pattern. Compared with average male failure rates from within-population crosses, the frequencies observed among individual France/Zambia cross directions ranged from no elevation at all (*i.e*. 0-2%) to as much as an order of magnitude greater (16.7%; Tables S4 & S5). Such divergent outcomes among crosses might not be expected under a model of numerous small effect BDMIs, in which case each cross might be expected to contain roughly comparable numbers of incompatibilities and rates of male failure. Instead, the striking variation in male failure rates among crosses may hint at the presence of large-effect interactions driving hybrid breakdown, with at least one interacting allele in each BDMI remaining polymorphic within its population. Further research will be needed to gain more detailed insights regarding the genetic architecture and molecular basis of these putative BDMIs.

Our finding that evidence of hybrid breakdown between recently-diverged populations is strongest for male reproduction follows theoretical predictions and previous observations that emergent reproductive isolation disproportionately affects the heterogametic sex, evidenced particularly between many *Drosophilae* (known commonly as Haldane’s Rule) (Haldane 1922; Coyne and Orr 1997; Orr 1997). Many common explanations of this trend invoke the X (or Z) chromosome, and a ‘large X effect’ has been described and confirmed in many model systems (Presgraves 2018). Our experimental design does not specifically address the role of the X chromosome in hybrid breakdown, since regardless of cross direction, F2 males receive their lone X chromosome from a mother who has one X from each parent. Hence, further genetic analysis will be needed to confirm the presence of X chromosome involvement in our observed male reproductive failure.

The variable effect of cross direction on male reproductive failure that we observed does imply a role for uniparentally-inherited factors, *i.e*. Y chromosomes, mitochondria, or other cytoplasmic factors such as maternally-inherited endosymbionts. We can not say whether the variability in cross direction effect is driven by within-population variation in the uniparental interactors or their partner loci (or both). Since both directional biases were observed among our set of France/Zambia crosses, it follows that at least two such factors would need to be involved in incompatibilities, *e.g*. both populations’ Y chromosomes, both populations’ mitochondria, or one population’s Y and mitochondria. In *Drosophila melanogaster*, polymorphism in both the mitochondria and Y chromosome have been shown to contribute to differential fitness of male offspring (Chippindale and Rice 2001), suggesting that possible factors contributing to reproductive failure here may indeed involve these genetic elements. In closely related species, Y chromosome variation has been suggested to contribute to hybrid sterility between *D. mauritania* and *D. simulans* (Bayes and Malik 2009). Alternatively, some models of early stage reproductive isolation also implicate mito-nuclear interactions (Ellison and Burton 2007, Rand *et al*. 2004). Backcross results in Table S6 do not allow inferences as to which uniparental element is driving reproductive failure in the current study, and both the mitochondria and Y chromosome remain interesting candidates for BDMIs between these populations.

Following up on the implication of uniparental elements in hybrid breakdown of male reproduction, we conducted backcrosses that, under a two-locus BDMI model, were predicted to yield much higher rates of male failure than seen among F2s. Instead, we observed the opposite pattern of much lower male failure among backcross offspring than among F2s (Figure 3). These results suggest that BDMIs involving uniparental elements do not reflect simple two-locus interactions, and specifically that one or more additional partners may include a recessive allele from the same source population as the uniparental element. Such an interaction is not possible to generate in our backcross design, as there are no scenarios where a homozygous autosomal or X-linked allele is present together with the uniparental chromosome being tested. This scenario is comparable to theoretical (Cutter 2012, Fraisse *et al*. 2014) and empirical studies (Phadnis 2011, Turner and Harr 2014, Phadnis *et al*. 2015) that suggest incompatibilities between isolated groups are likely to be polymorphic and to involve multiple partner loci. The implication of multigenic incompatibilities is consistent with the patterns of ancestry-mediated epistatic selection detected by an analysis of African and European ancestry in an admixed North American population, which found hub-like patterns of “ancestry disequilibrium” that could potentially be explained by complex incompatibilities (Pool 2015).

The existence of hybrid breakdown between populations that diverged within the last 13,000 years (Sprengelmeyer *et al*. 2020) implies that incompatible variants were either already present at the time that populations split, or else they arose and increased in frequency rapidly thereafter. This early emergence of hybrid breakdown is intriguing in light of a study of isolation between multiple *Drosophila* species which found that postzygotic isolation such as hybrid sterility usually evolves more slowly than prezygotic forms of isolation (Turissini *et al*. 2018). We note that some level of prezygotic isolation, in the form of a polymorphic unidirectional mating preference, does also exist between these populations (see Introduction). Hence, results from this and prior studies in *D. melanogaster* emphasize that both pre- and post-zygotic isolation can begin to emerge in the earliest stages of population divergence.

Evidence for hybrid breakdown impacting male reproduction was strongest between populations from France and Zambia. An additional possible signature of elevated male reproductive failure was also observed between France and Cameroon hybrids, but not explored further in the present study. The involvement of France in both of these pairings is interesting in that this population has the most distinct allele frequencies among the four studied, as a consequence of the out-of-Africa bottleneck (Lack *et al*. 2016b, Sprengelmeyer *et al*. 2020). However, no strong enrichment of hybrid male failure was observed between France and Ethiopia (both of which have adapted to colder environments; Pool *et al*. 2017), even though these populations have the highest *FST* of any population pair in our study (0.312; Lack *et al*. 2016b). Instead, France shows potential signals of hybrid breakdown with the two warm-adapted African populations in our study (Figure 1D). Our results thus far can not suggest to what degree incompatible variants have increased in frequency due to adaptive evolution (Schluter and Conte 2009) versus genetic drift (Coyne and Orr 2004, Schiffman and Ralph 2021). Still, it is worth noting that incomplete sweeps appear to be common in *D.melanogaster* (Garud and Petrov 2016; Vy and Kim 2017; da Silva Ribeiro *et al*. 2022) and to underlie cases of locally adaptive trait evolution (Bastide *et al*. 2016; Sprengelmeyer and Pool 2021; Sprengelmeyer *et al*. 2022). Hence, the polymorphic nature of hybrid breakdown that we observe is consistent with adaptive as well as neutral hypotheses. If the genes underlying these apparent BDMIs can ultimately be identified, population genetic patterns at these causative loci may inform this important question. Future studies could also choose to further quantify hybrid breakdown between some population pairs that have high genetic differentiation and low environmental differentiation, and other pairs with the opposite relationships, in order to provide further clues as to whether genetic drift or environmental adaptation is more important in driving early stage reproductive isolation.

### Conclusions

Early stages of incomplete reproductive isolation may include hybrid breakdown, when recessive incompatibilities that are masked in the first generation of interbreeding are exposed in later generation hybrids. Here, we provide strong evidence for the existence of hybrid breakdown in male reproductive performance (but no clear evidence for female reproduction or viability) between recently-diverged populations of *D. melanogaster*. We find that the rate of reproductive failure among males from crosses between France and Zambia populations is significantly greater than within population crosses. The apparent cross and cross-direction variability that exists among these crosses suggests that incompatibilities are polymorphic and involve uniparental factors in some cases. Further, backcross results provide further evidence of a complex basis of hybrid breakdown, implying the existence of multi-locus incompatibilities. Taken together, our results suggest that incompatibilities have arisen rapidly after populations diverged, that they remain variable, and that at least some of them have a complex genetic architecture. In light of the experimental tractability of *D. melanogaster*, the genetic tools available, and our knowledge of this species’ genome and its diversity, the existence of hybrid breakdown between *D. melanogaster* populations appears to offer promising opportunities for future investigations of the molecular basis of hybrid breakdown in early stage reproductive isolation.

## ACKNOWLEDGEMENTS

This research was supported by NIH grants R35 GM13630 (to JEP), T32 GM007133, and T32 HG002760. We thank members of the Pool lab for helpful comments on draft versions of this manuscript.

## AUTHOR CONTRIBUTIONS

MJL and JEP designed the research, MJL, CED, and TJB performed the research, MJL and JEP wrote the paper.

## DATA ACCESSIBILITY

All data produced for this study is reflected in the supplementary tables.

